# Variation in the response to antibiotics and life-history across the major *Pseudomonas aeruginosa* clone type (mPact) panel

**DOI:** 10.1101/2024.01.15.575732

**Authors:** Leif Tueffers, Aditi Batra, Johannes Zimmermann, João Botelho, Florian Buchholz, Junqi Liao, Nicolás Mendoza Mejía, Antje Munder, Jens Klockgether, Burkhard Tümmler, Jan Rupp, Hinrich Schulenburg

## Abstract

*Pseudomonas aeruginosa* is a ubiquitous, opportunistic human pathogen. Since it often expresses multidrug resistance, it is ranked by the World Health Organization among the top 3 high priority pathogens, for which new treatment options are urgently required. An evaluation of new treatments is usually performed experimentally with one of the canonical laboratory strains (e.g., PAO1 or PA14). However, these two strains are unlikely representative of the strains infecting patients, because they have adapted to laboratory conditions and do not capture the enormous genomic diversity of the species. Here, we characterized the major *P. aeruginosa* clone type (mPact) panel. This panel consists of 20 strains, which reflect the genomic diversity of the species, cover all major clone types, and have both patient and environmental origins. We found significant strain variation in distinct responses towards antibiotics and general growth characteristics. Only few of the measured traits are related, and if so, only for specific antibiotics. Moreover, high levels of resistance were only identified for clinical mPact isolates and could be linked to known AMR (antimicrobial resistance) genes in the sequenced genomes. One strain also produced highly unstable AMR, indicating an evolutionary cost to resistance expression. By linking isolation source, growth, and virulence to life history traits, we further identified specific adaptive strategies for individual mPact strains towards either host processes or degradation pathways. Overall, the mPact panel provides a reasonably sized set of distinct strains, enabling in-depth analysis of new treatment designs or evolutionary dynamics in consideration of the species’ genomic diversity.

**Importance:** New treatment strategies are urgently needed for high risk pathogens such as the opportunistic and often multidrug resistant pathogen *Pseudomonas aeruginosa*. Here, we characterize the major *P. aeruginosa* clone type (mPact) panel. It consists of 20 strains with different origins that cover the major clone types of the species as well as its genomic diversity. This mPact panel shows significant variation in (i) resistance against distinct antibiotics, including several last resort antibiotics, (ii) related traits associated with the response to antibiotics, and (iii) general growth characteristics. We further developed a novel approach that integrates information on resistance, growth, virulence, and life-history characteristics, allowing us to demonstrate the presence of distinct adaptive strategies of the strains that focus either on host interaction or resource processing. In conclusion, the mPact panel provides a manageable number of representative strains for this important pathogen for further in-depth analyses of treatment options and evolutionary dynamics.

## Introduction

Evolving human pathogens are a major global challenge for human wellbeing. The recent COVID-19 pandemic has been a dramatic reminder of this challenge. Bacterial pathogens pose a similar threat, because of their ongoing adaptation to the human host and particularly the emergence and spread of antimicrobial resistance (AMR). A recent worldwide analysis revealed that approximately 5 million deaths per year are associated with infections caused by AMR-expressing pathogens, and of these, a total of 1.27 million deaths per year can be directly attributed to AMR (1). AMR has thus been declared as one of the main areas requiring urgent action by the World Health Organization (2). The Gram-negative bacterium *Pseudomonas aeruginosa* is an example of a top priority pathogen, for which new treatment options are urgently needed (3) and which is part of the highly problematic group of the ESKAPE pathogens (4, 5). This bacterium is a ubiquitous opportunistic human pathogen that is found in diverse environments and can cause various infections in humans, often acquired in a hospital context (6). Frequent are *P. aeruginosa* infections of wounds or of the lung, the latter often chronic in patients with cystic fibrosis (CF), bronchiectasis or chronic obstructive pulmonary disease (COPD) (6). This pathogen commonly expresses AMR, often against multiple antibiotics, making it difficult and in some cases impossible to treat.

Due to its medical importance, *P. aeruginosa* has become a model for studying the mechanistic basis as well as the evolution of AMR. A particular focus is on understanding the genetics of AMR. This pathogen is notorious for having a large as well as flexible genome with diverse AMR genes, often found on mobile genetic elements, especially integrative and conjugative/mobilizable elements (ICEs/IMEs) and also plasmids (7, 8). Genes responsible for AMR against diverse antibiotics have been identified, including widely used antibiotics like those from the group of β-lactams (9, 10) as well as reserve antibiotics, such as colistin (11), or newly developed drugs, such as ceftolozane (12). This bacterium is further able to express phenotypic resistance, which may be based on forming biofilms, producing bacterial persisters, or through the expression of heteroresistance (13). These phenotypic resistance mechanisms often complicate diagnostics and make it difficult to predict AMR from the genome sequence alone (14–16). Moreover, the majority of the functional analyses use two main laboratory strains, PAO1 and PA14 (17–19). Thus, we often lack information on whether the identified mechanisms underlying AMR in these two laboratory strains apply to the clinically relevant pathogen strains and can assist the improvement of diagnostics as well as treatment designs.

The aims of the current study are to characterize a panel of *P. aeruginosa* strains, which are representative of the genomic diversity encountered in this species and which are still of a manageable size for in-depth functional as well as evolutionary analyses. It builds on the strains which have been isolated previously by the ‘Pseudomonas Genomics’ group at Hannover Medical School and includes the major *P. aeruginosa* clone types (mPact) (20–22). These mPact strains have been isolated from different environments as well as different types of human infections, and vary in virulence towards distinct hosts (22, 23). In the current study, we describe these strains and their genomic distribution. We further characterize variation in AMR in detail using distinct, commonly applied approaches, including diagnostic automated susceptibility testing (AST), Etests, and disc diffusion assays, in consideration of a range of clinically relevant antibiotics. We also examine the presence of microcolonies and variation in the shape of time-kill curves as indicators of additional defense responses to antibiotics, and relate these characteristics to variation in general growth parameters (i.e., lag time, exponential growth rate, yield). In addition, we reconstructed metabolic models and predicted life history traits that could predict growth and clinical parameters. These characterizations yield new insights into the variation in AMR and related responses in a representative set of strains for *P. aeruginosa* and thereby provide a reference for future in-depth molecular as well as evolutionary analysis of AMR or virulence in the mPact panel. The mPact panel is available to the scientific community as a resource for future research.

## Results

### The major *P. aeruginosa* clone type (mPact) strain panel covers a large part of the genomic diversity of the species

We focused our work on the strain panel originally selected to represent the interclonal diversity of the major clones in the present-day *P. aeruginosa* population (20, 22). Using our recent whole genome sequence analysis, we can show that the mPact panel is distributed across a large part of the species’ phylogeny (Fig. 1, Table 1). It includes members of all three main phylogroups. Clinical and environmental mPact strains are available for both main phylogroups A and B. Some strains are more closely related to the two main reference strains PA14 (mPact strain H02) and PAO1 (mPact strain H11), while the remaining strains are placed in distinct phylogenetic groups. Genome size varied among mPact strains, ranging from 6.26 Mb to 7.27 Mb (Table 1). The larger genomes generally harbored more mobile genetic elements. In detail, three strains contain one plasmid, while one other strain has three plasmids. Up to six integrative conjugative elements (ICEs) are found in the mPact genomes. Overall, the wide phylogenetic distribution of the included strains and their origin from both clinical and environmental samples makes the mPact panel a small, representative sample for in-depth analysis.

**Figure 1.**
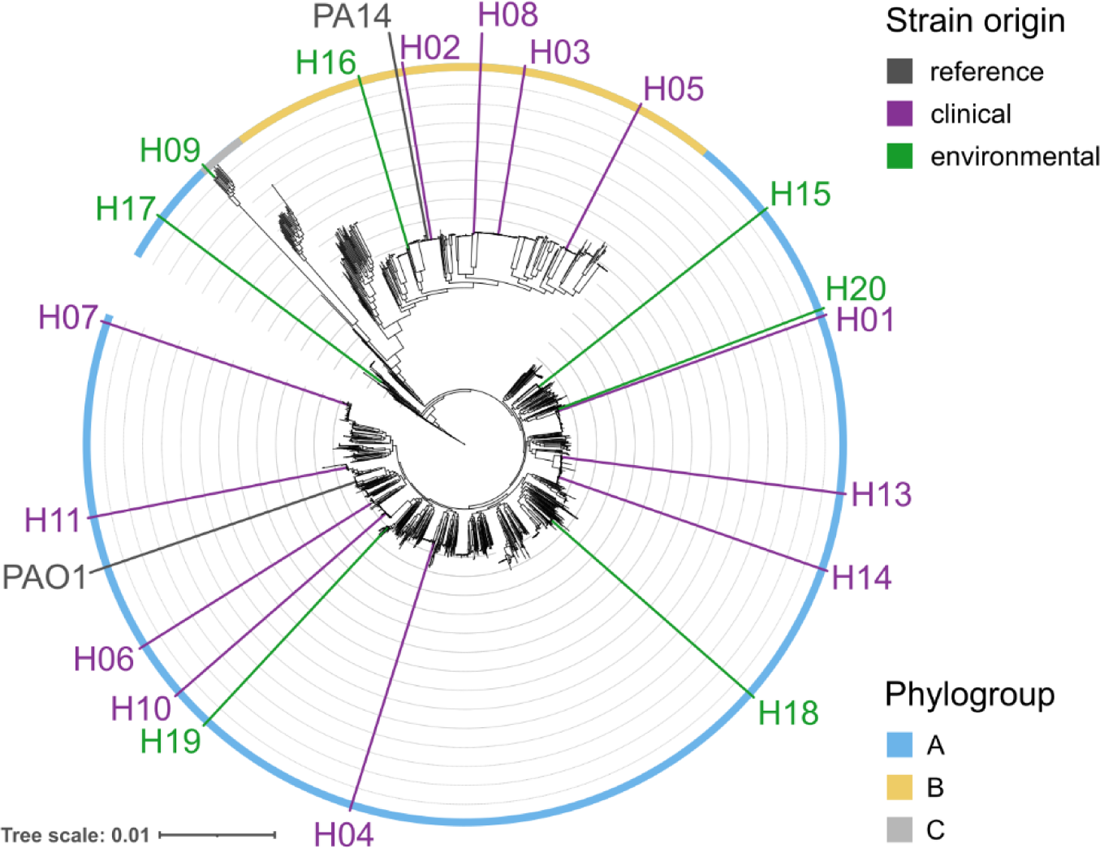
Distribution of the mPact strains across the genome-sequence-inferred phylogeny of *P. aeruginosa*. Maximum likelihood of the softcore-genome alignment of 2009 *Pseudomonas aeruginosa* isolates. The scale bar indicates genetic distance. Outside colored ring represents phylogroup placement within the species pangenome. The mPAct panel strains are indicated and placed according to the phylogenetic position; their origin is denoted by color. Figure modified from Botelho et al. (8) and used with permission of Elsevier/eBioMedicine.

**Table 1.**
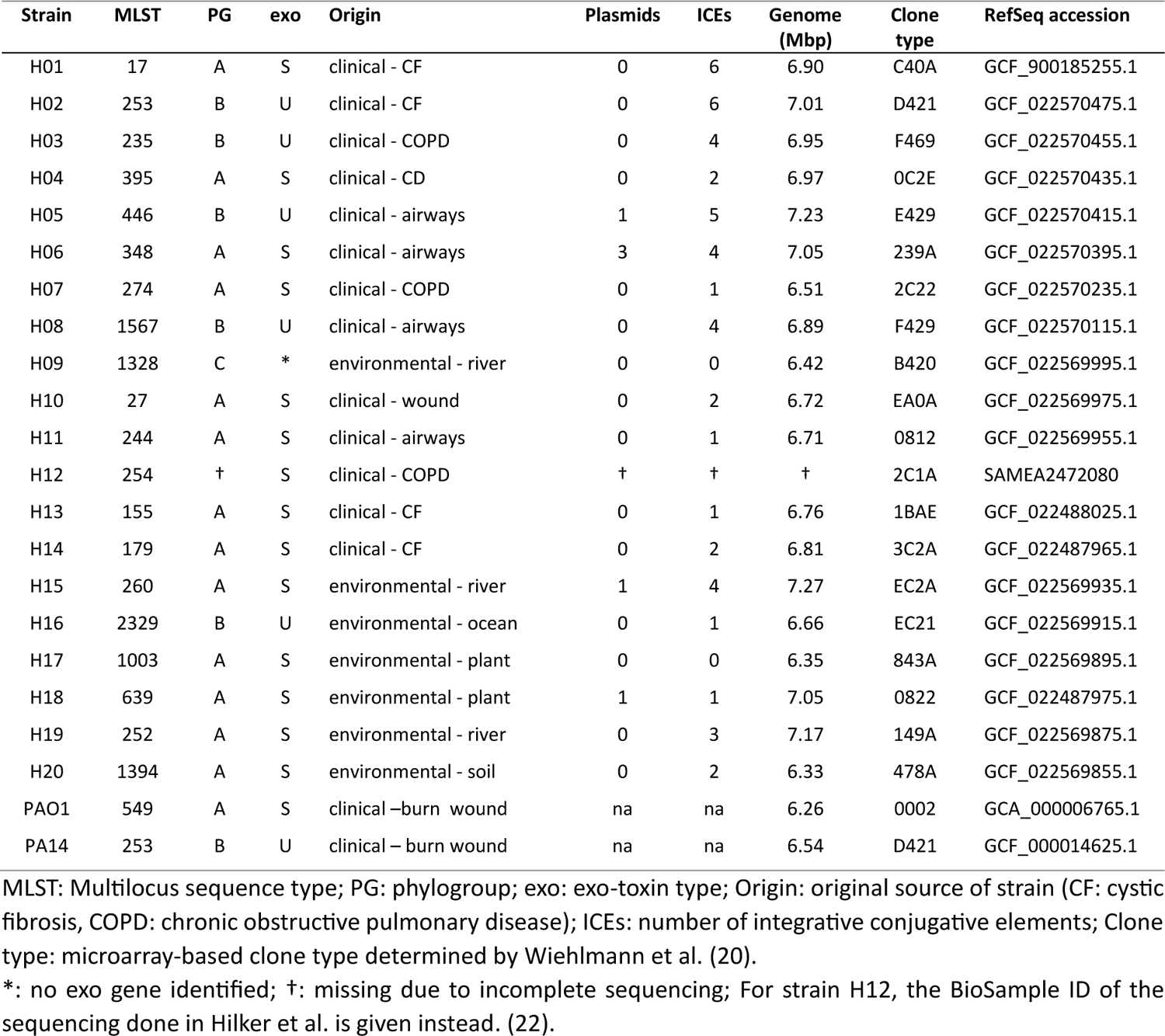
General information on the origin and phylogenomic characteristics of the mPact panel.

### Variation in antibiotic resistance across mPact strains is related to origin but not phylogroup

Antimicrobial resistance varied substantially across the panel strains (Figure 2A). Using EUCAST resistance breakpoints, six of 22 strains were classified as clinically resistant against at least one substance. Resistance against ciprofloxacin was most frequent (6 cases), while resistance to carbapenems was rare. All strains exhibiting resistance were clinical in origin, while the environmental strains showed only slight variation in overall high drug susceptibility. No pattern of resistance matching phylogroup distribution was apparent. All strains showed low MICs against the last resort drugs ceftazidime/avibactam, ceftolozane/tazobactam, cefiderocol, and also colistin. Resistance measurements using gradient strip and disc diffusion methods on M9 minimal agar yielded similar results, but did not confirm resistance to the beta-lactam drugs piperacillin and ceftazidime (Figure S1). The quantitative results of disc diffusion and gradient strip MICs generally covaried well (Figure 1B), with some restrictions in this strain set for colistin and ceftolozane/tazobactam.

**Figure 2.**
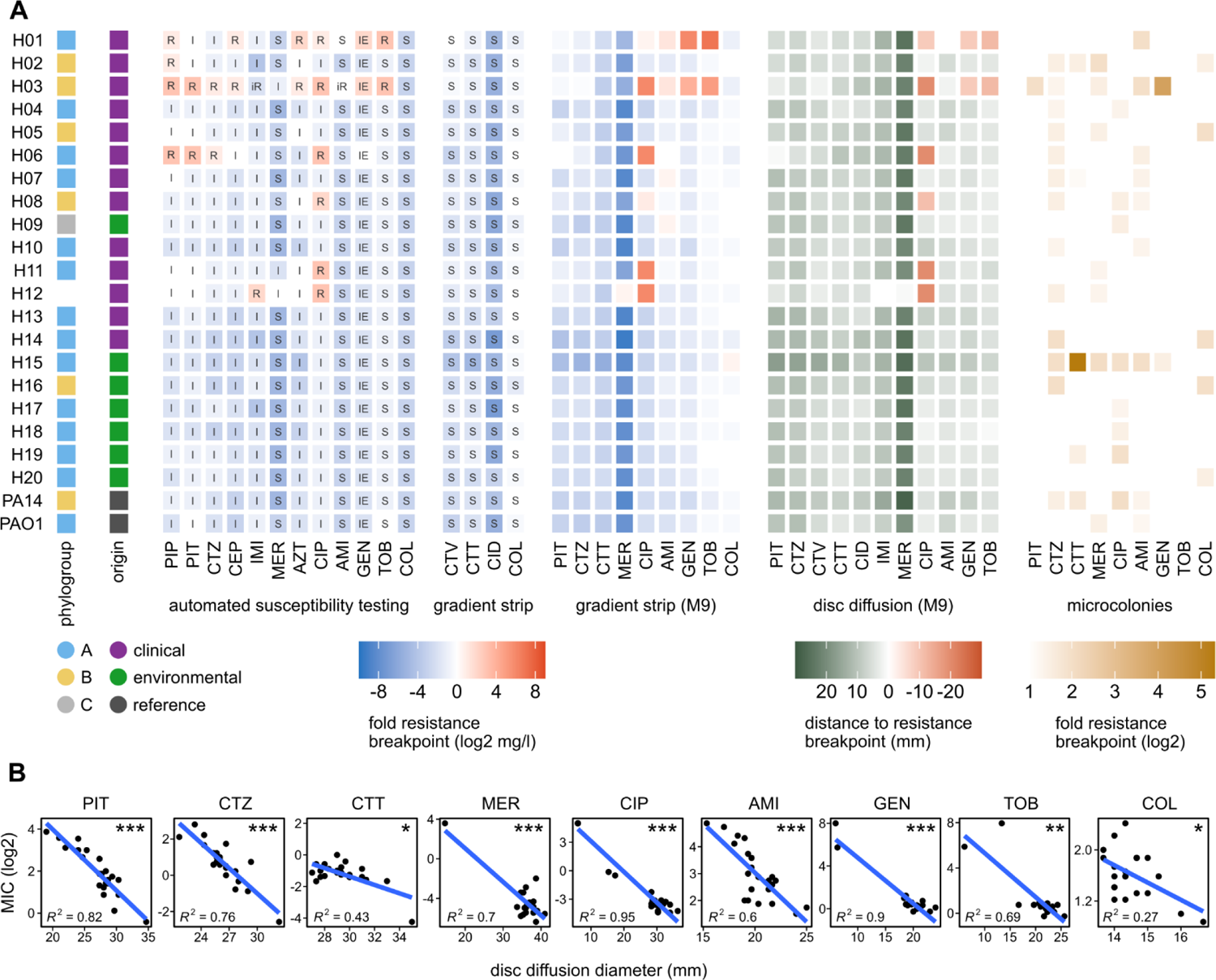
Variation in resistance across the mPact panel. (A) Resistance measurements in (from left to right) automated susceptibility testing (AST, VITEK2), reserve antibiotics on Mueller-Hinton agar (gradient strip, Etest) and Etest on M9 minimal medium agar, displayed as their fold ratio of minimum inhibitory concentration (MIC) to EUCAST resistance breakpoint; disc diffusion on M9 minimal medium agar, displayed as zone of inhibition diameter minus the EUCAST resistance breakpoints in mm; and effect of microcolonies on M9 Etest resistance data. Antibiotic abbreviations: PIP = piperacillin, PIT = piperacillin + tazobactam, CTZ = ceftazidime, CEP = cefepime, IMI = imipenem, MER = meropenem, AZT = aztreonam, CIP = ciprofloxacin, AMI = amikacin, GEN, gentamicin, TOB = tobramycin, COL = colistin, CTV = ceftazidime + avibactam, CTT = ceftolozane + tazobactam, CID = cefiderocol. Box labels in the heatmaps indicate clinical resistance classification according to EUCAST, where appropriate: S = susceptible, I = susceptible at increased exposure, R = resistant, iR = inferred resistance from related substances despite a minimal inhibitory concentration below the breakpoint. IE: insufficient evidence for clinical use. (B) Regression of M9 gradient strip MIC and M9 disc diffusion diameter. R^2^ is the coefficient of determination of the linear regression model. Symbols in the upper right corners indicate the p-values of the t-test of R^2^ = 0, with * = *p* < 0.05, ** = *p* < 0.01, *** = *p* < 0.001.

**Figure 3:**
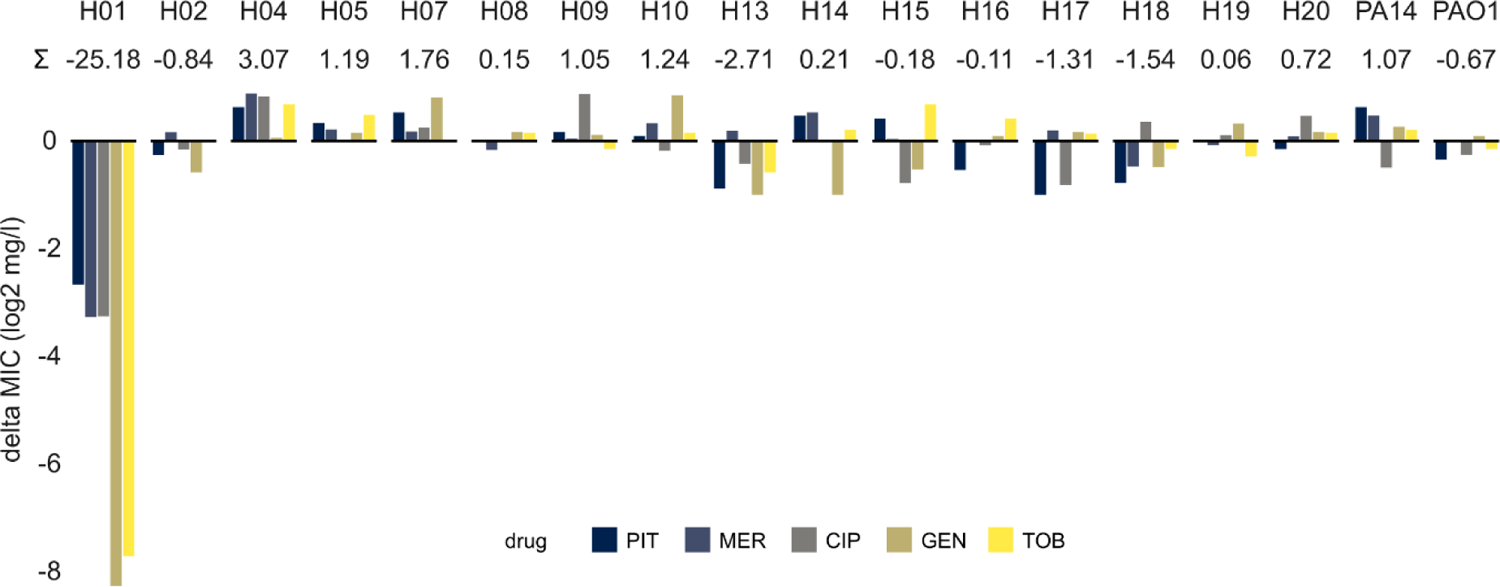
Stability of antibiotic resistance. The MICs of five drugs (for abbreviations, see legend of figure 2) were measured after fourteen days of evolution on drug-free medium. Ancestral MICs were subtracted from the evolved MIC, so that resistance loss results in negative delta MIC values. The sum of resistance change (log2 transformed) is given beneath the strain names. Strains with a high ancestral MIC to ciprofloxacin were excluded.

Microcolony formation inside zones of inhibition of the agar-based methods was assessed to find indications of monoclonal heteroresistance, a phenomenon in which a genetically clonal bacterial population exhibits a range of MICs against an antibiotic in the form of individual colonies growing inside the inhibition zones. Microcolonies were observed in numerous strain/drug combinations (Figure 2A, rightmost panel), but mainly resulted in a less than twofold MIC increase, with the notable exception of H15 and ceftolozane/tazobactam, where colonies resulted in a 32-fold MIC increase. Some drugs caused little to no microcolony formation (piperacillin/tazobactam, tobramycin), while it was frequent in others, especially ceftazidime. Similarly, some strains exhibited almost no microcolony formation (H13, H20), while others reacted to many drugs in this manner (H15). Overall, possible tendencies towards heteroresistance could be observed. Strong MIC increases due to microcolonies could be observed in two cases.

Resistance phenotypes were tested for their stability against spontaneous loss by repeatedly culturing individual strains in drug-free medium. While the majority of resistances remained stable and fluctuated only mildly around the ancestral MICs, strain H01 showed a dramatic loss of all tested resistances. This loss resulted in an MIC change to one quarter of the ancestral value for piperacillin/tazobactam, meropenem and ciprofloxacin, and an even more pronounced loss of aminoglycoside resistance. This strong effect hints at a strain-specific configuration of resistance mechanisms with unstable expression or high fitness cost in the absence of selective pressure.

### Substantial variation in time-kill dynamics among mPact strains towards high concentrations of distinct antibiotics

The kinetics of bacteria-antibiotic interactions over time can give indications of a bacterial strategy for the survival of lethal drug doses, which is distinct from phenotypic resistance. A slow decrease in cell density after addition of a drug despite above-MIC concentrations can hint at the presence of a tolerant phenotype, in which survival is based on a decrease in cellular activity in order to avoid or repair the cellular effects of the drug, rather than the expression of a resistance phenotype. We created time-kill curves of the panel strains against four distinct antibiotics at drug concentrations at least 32-fold the strain MICs and quantified the populations’ responses by calculating the area under the time-kill curve (AUC) (Figure 4A). Strong differences between the antibiotics were observed, with ciprofloxacin and gentamicin resulting in rapid killing and thus small AUCs, while the beta-lactam antibiotics ceftazidime and meropenem killed populations more slowly and sometimes not completely.

**Figure 4:**
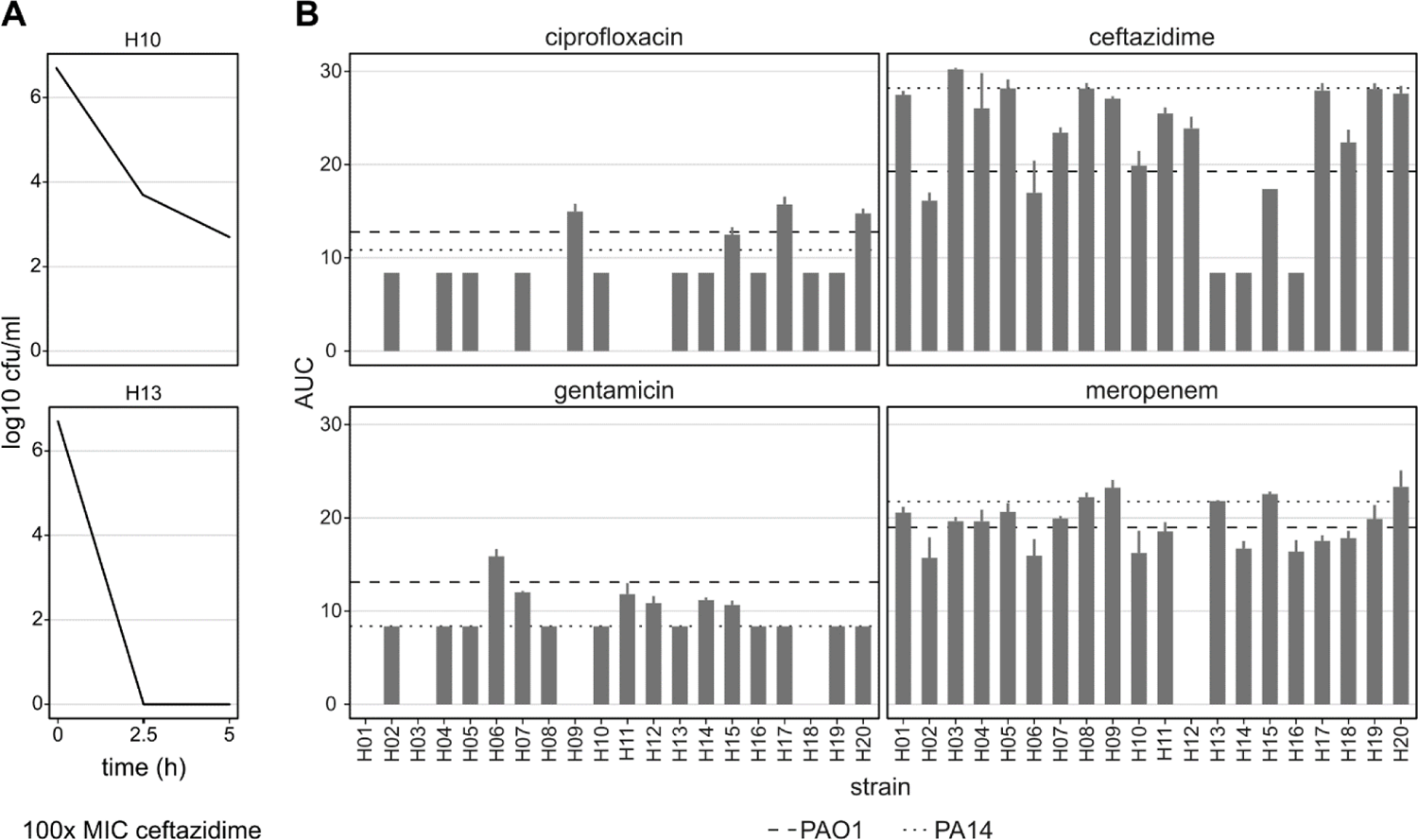
Variation in time-kill response among mPact strains towards distinct antibiotics. (A) Illustration of test principle. Strains were exposed to drug concentrations at least 32x above the respective MICs and cell numbers counted after 2.5 and 5 hours. The area under the time-kill curve (AUC) is used to compare strains. (B) AUCs per strain, split by test antibiotic. Missing bars indicate combinations that were not tested due to high strain MICs. The dotted lines indicate the respective average values for the laboratory strain PA14 and dashed lines those for the laboratory strain PAO1. Error bars represent standard deviation of three replicates.

None of the examined strains exhibited a clear increase in AUC against all drugs, but some cases of variable response were observed. For example, strain H06 showed above average AUC against gentamicin, but below average AUC against the beta-lactams, indicating drug-specific responses. No correlations between strain MIC and AUC were observed, with the exception of ceftazidime (Figure S2), in which three strains showed very rapid killing. These strains had low, but not the lowest ceftazidime MICs. Because this response to antibiotics could also be linked to overall bacterial growth characteristics, we next examined growth in the absence of drugs.

### mPact strains vary in general growth characteristics

Several panel strains grew rapidly in M9 minimal medium (Figure 5, Supplementary Table S1), with short lag times and high growth rates (H05, 07, 10, 11, 12, 13). In contrast, some strains were markedly slow (H01, H15), with opposing parameters. This indicates a link between these main growth characteristics, which in combination results in matching differences in area under the growth curve. However, this trend was not universal. H14, for example, showed a relatively low growth rate, but a very short lag phase, resulting in an overall large AUC. Optical density-based carrying capacity, which has an increased effect on AUC the longer the experiment runs, also varied significantly.

**Figure 5.**
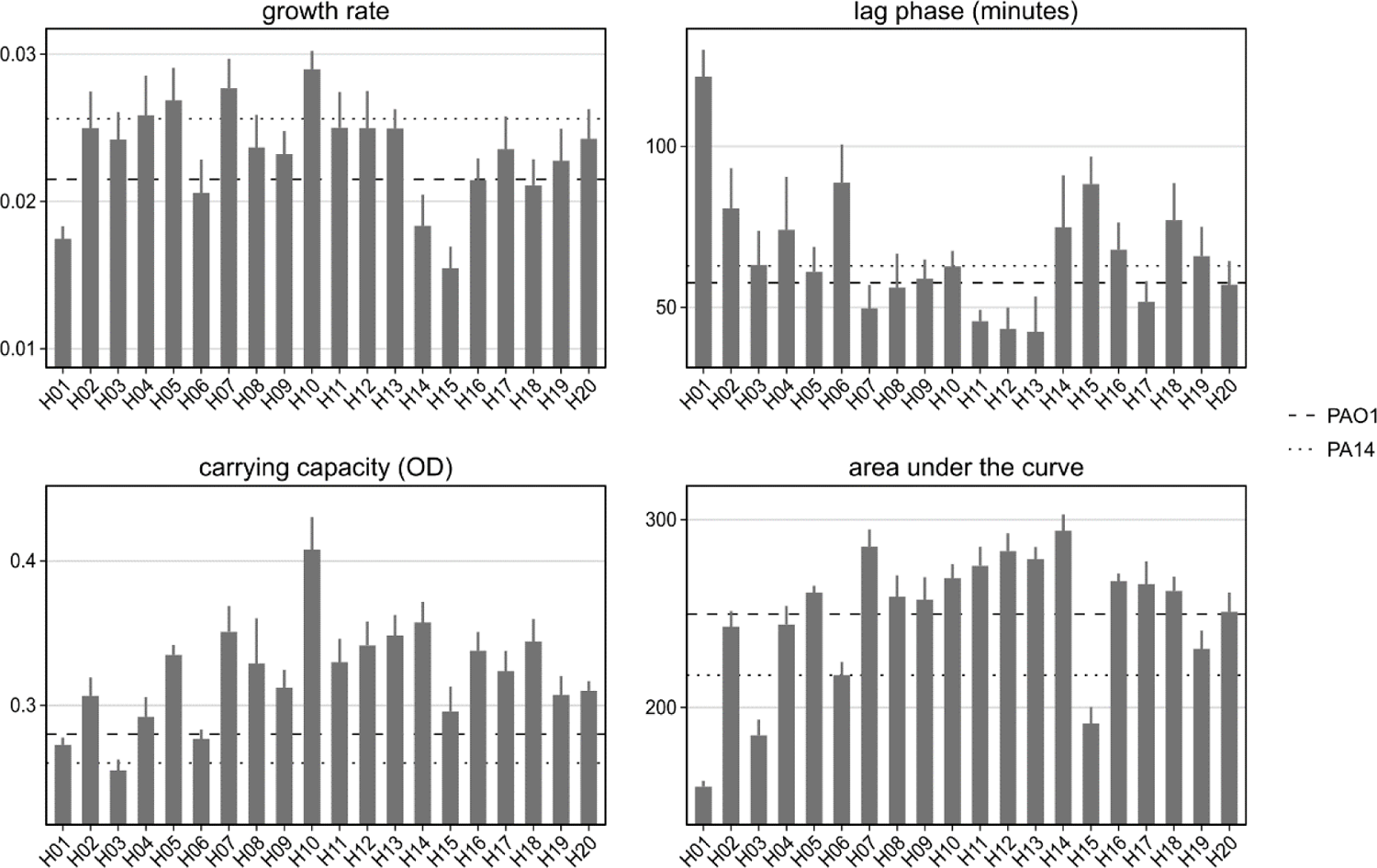
Variation in growth characteristics across mPact strains. Bacterial growth was characterized in the absence of antibiotics in M9 medium. The characteristics of bacterial growth were assessed as (i) the growth rate (top left panel) and (ii) lag time (top right panel), inferred from a log-linear growth model (24) using the growthrates package in R, (iii) carrying capacity (bottom left panel), recorded as optical density (OD) at the end of the growth season, and (iv) area under the curve (AUC) of OD measurements across the entire growth period. The dashed and dotted lines indicate the average values for the laboratory strains PAO1 and PA14, respectively. Error bars represent standard deviation of at least 8 replicates. Further details are provided in Supplementary Table S1.

We tested the observed variation of growth parameters against a number of strain characteristics and phenotypes. For a cumulative measurement of strain resistance to use in these analyses, we tested multiple aggregate scores. On the one hand, the sum of disc diffusion diameters resulted in a symmetrical distribution with a mean value of 301 mm (Figure S3, purple curve). Sorting the strains by this score confirmed that the most resistant strains were of clinical origin, even though some environmental strains also showed a certain level of resistance (Figure S4). In addition, calculating the sum of gradient test MICs yielded a strongly skewed distribution with two major outliers (Figure S3, cyan curve), which persisted after transforming the data as relatives to the drug-specific breakpoints and log2-transforming the MICs to compensate for the doubling dilution series of MIC measurements (Figure S3, orange curve). Neither of these cumulative resistance scores covaried significantly with the growth parameters (Figures S5 and S6). No significance test was performed for the MIC-based resistance score, due to H01 being a strong outlier. This was due to the aminoglycoside MICs in strain H01 being extremely high, while disc diffusion diameters were closer to those of the other strains, indicating a poor correlation of these methods for very high aminoglycoside concentrations.

### Resistance phenotypes can be related to resistance genes in the sequenced genomes

The genetic makeup of the strains was tested for contribution to the phenotypes described above. We found no significant correlations between phylogroup, origin, number of plasmids and number of ICEs on the one hand, and cumulative resistance and growth characteristics on the other, with one exception. A higher number of ICEs correlated with a longer lag phase (Figure S7), possibly indicating a growth cost of these mobile elements. All tested correlations are summarized in Supplementary Table S2.

Finally, we tested the genomes for known resistance determinants to explain the variation in phenotypes. AMRfinder queries yielded a number of resistance genes (Figure 6, Supplementary Table S3), which matched the observed resistances. In detail, strains with aminoglycoside resistance carried strain-specific aminoglycoside-modifying enzymes (i.e., ant-(2’’)-Ia in H01, aadA6 in H06, and aph(3’)-IIa, aph(6)-Ic and aac(6’)Ib4 in H03). Point mutations in quinolone-resistance determining regions (QRDR) gyrA and parC were found in strains with ciprofloxacin resistance. AMRfinder also found the sulfonamide resistant variant of dihydropteroate synthase (sul1) and the antiseptic resistance conferring variant of qacE (qacEdelta1). Neither of these phenotypes were tested in our study, but were found in highly resistant clinical isolates, indicating further adaptations to a hospital environment. No clear beta-lactam resistance genes were identified. We therefore performed a manual search of the known beta-lactam resistance genes blaPDC, ampD, ampDh3, ampR, dacB, ftsI and oprD. Genetic distance in their amino acid sequences in some cases partially matched the pangenome phylogroups (ampDh3, oprD), but was not associated with beta-lactam resistance except in one case (Figure S8). We identified a frameshift mutation in dacB in strain H06. Loss of function of this gene, which likely results from this mutation, is a known beta-lactam resistance mechanism in *P. aeruginosa*, matching this strains’ elevated MICs against piperacillin and ceftazidime. No confirmed resistance mechanism was found in H03, which showed beta-lactam resistance in the automated susceptibility testing only (Figure S6). It should be noted that the only strain with distinct reduction in carbapenem susceptibility was not included in this analysis due to an incomplete genome. All in all, the identified resistance genes explain the majority of observed resistance.

**Figure 6.**
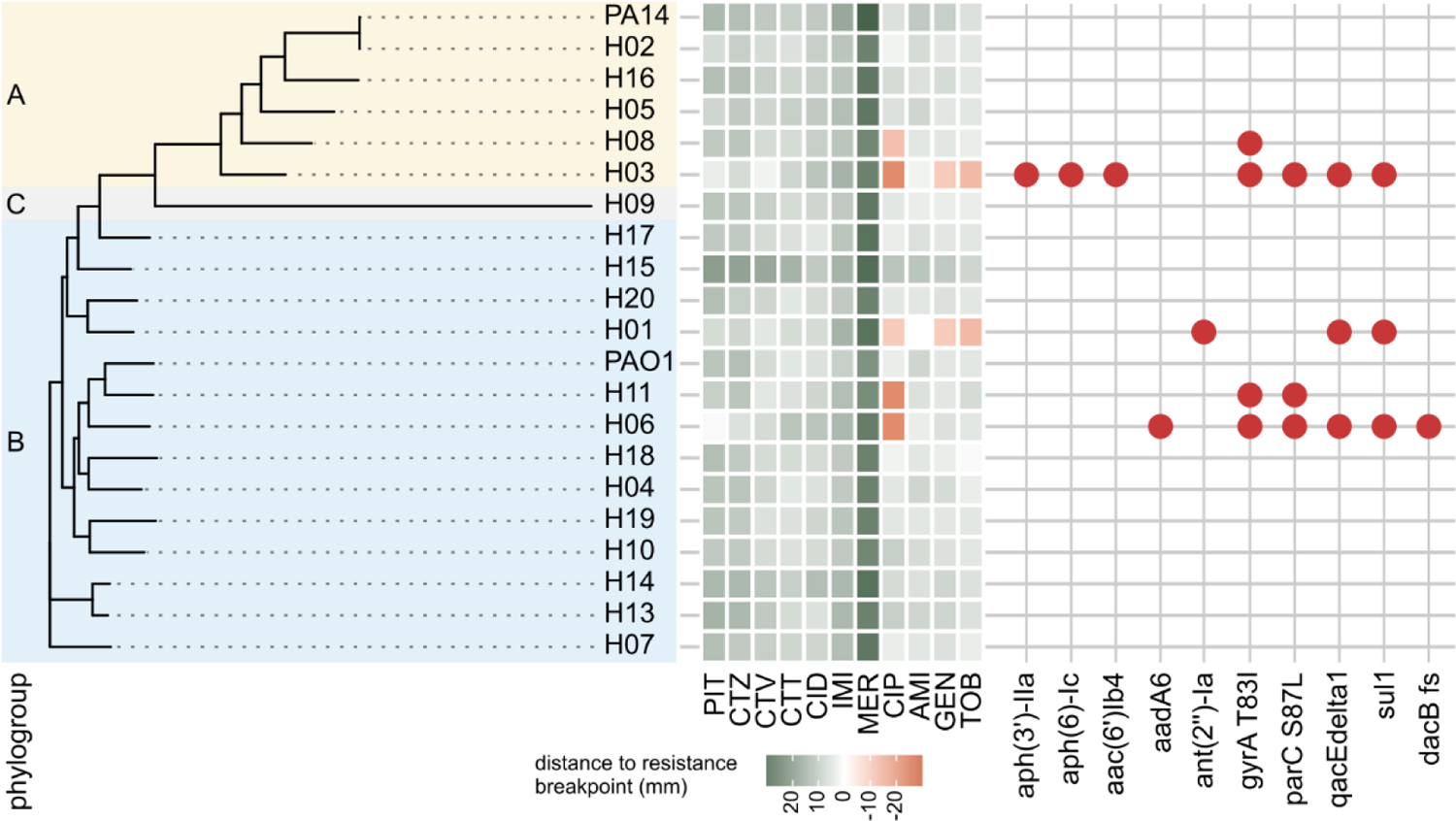
Genomic resistance determinants are associated with resistance phenotypes. Strains are arranged according to their core genome phylogeny and shaded by phylogroup (left part). The heatmap indicates strain resistance to the given drugs plotted as the distance of disc diffusion diameter to the drug resistance breakpoint, resulting in resistant values given as negative values (heatmap as in Fig. 2A). Resistance gene hits from AMRfinder are presented in the grid on the right. A frameshift in dacB was additionally identified in one strain through amino acid sequence alignment. Antibiotics abbreviations see legend of figure 2. Details of identified genes see text. Please note that a sixth strain with resistance (strain H12) is not included here because of incomplete genome sequence information, thus precluding inference of AMR genes in its genome.

### Genome sequence analysis predicts life history characteristics, growth and virulence phenotypes

We further explored the genomic capacity of the mPact panel by reconstructing metabolic models using a recently introduced pipeline called Bactabolize that especially accounts for differences between strains (Supplementary information). While we did not find variations when predicting growth rates by flux balance analysis, the metabolic models differed in their amount of metabolic reactions. Therefore, we performed an enrichment analysis to check for differences between strain groups. We found an enrichment of reactions involved in carbohydrate, amino acid, and secondary metabolite metabolism for environmentally obtained strains (Figure 7E). Next, we inferred microbial life history traits from genomes using GO terms, pathways, subsystems, secondary metabolites, carbohydrate-active enzymes, and virulence genes (see Methods; supplementary information). We identified traits that could predict microbial phenotypes in a multivariate analysis. In detail, the source of isolation (environment or clinic) could be predicted with an accuracy of 0.95 (determined by leave-on-out cross-validation): the capacity for posttranslational modification and osmotic stress response was indicative for clinical isolates, whereas lipid metabolism, carbohydrate degradation, and CRISPR genes were characteristic of environmental isolates (Figure 7A). Interestingly, we found contrasting directions in amine and polyamine metabolism with degradation in environmental and biosynthesis in clinical isolates (Figure 7A). Similarly, the virulence of strains in a mouse model (high, moderate, low) was estimated with an accuracy of 0.74 by the number of effector delivery systems (high virulence) and carbohydrate degradation pathways (low virulence; Figure 7B). Moreover, high growth rates were associated with host interaction processes and heat response, whereas slow growth was linked to cell cycle processes and the number of pseudogenes (Figure 7C). Compared to the experimental data, the predicted traits captured the growth variance accurately (Spearman correlation R=0.86, p-value<0.01; Figure 7D). In summary, life history analysis suggested partitioning of mPact strains into environmentally-derived, slower-growing carbohydrate degraders and clinically derived, host-associated, stress-tolerating, fast-growing pathogens.

**Figure 7.**
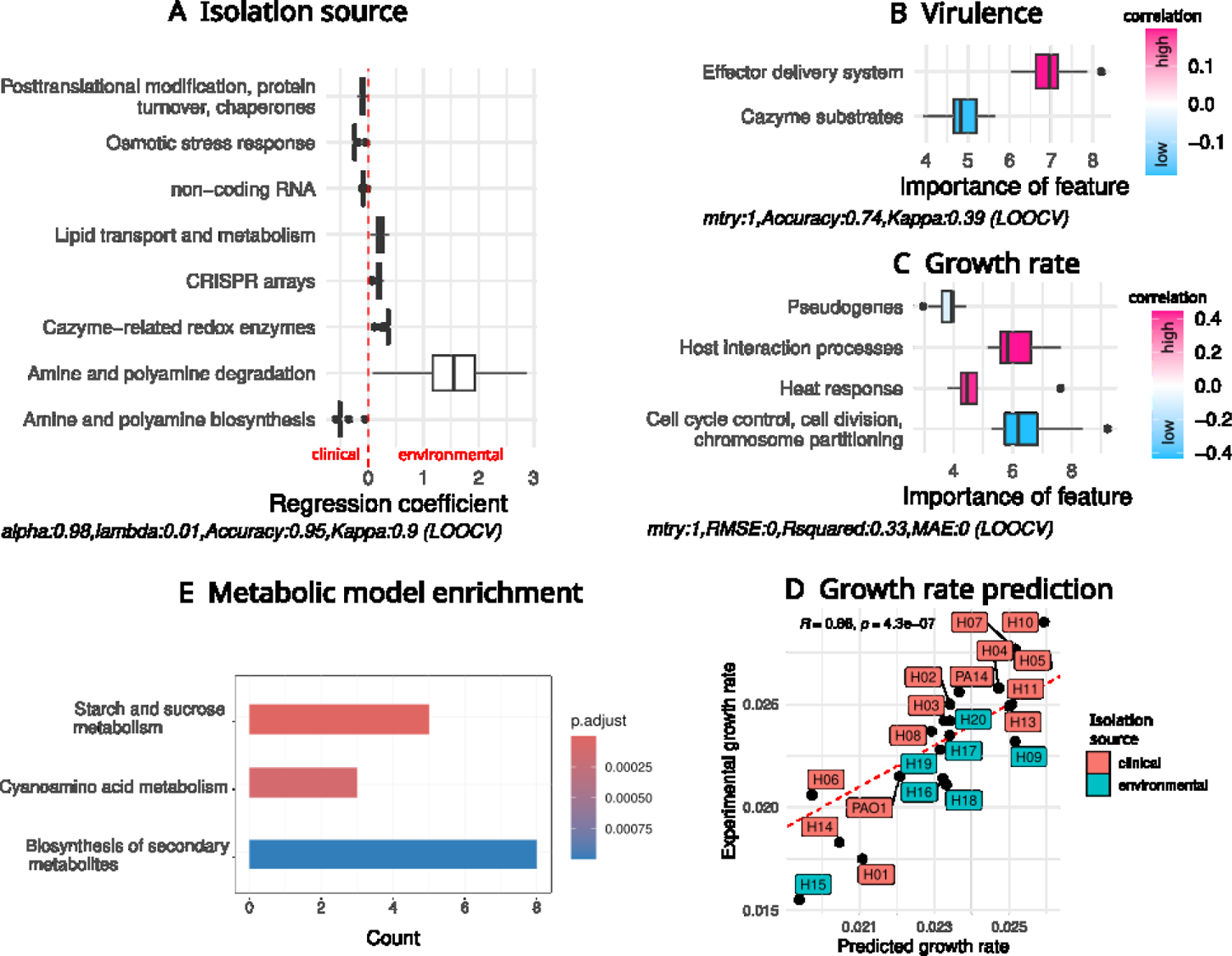
Genome sequence analysis and metabolic models predict life history characteristics of mPact strains. (A) Prediction of strain isolation source (environment or clinic) by life history traits using a lasso classifier. Accuracy was determined by leave-one-out-cross-validation (LOOV). (B) Inference of virulence (high, moderate, low) of mPact strains by life history traits using random forest classifier. Accuracy was determined by LOOV. (C) Prediction of strain maximal growth rates by life history traits using random forest regression. Root-mean-square error (RMSE) and *R^2^* were determined by LOOV. (D) Correlation of experimental maximal growth rates and the growth rates predicted by the random forest regression model from (C). (E) Enrichment of metabolic pathways found in metabolic models that were isolated from the environment compared to clinical isolates.

## Discussion

Our study identified substantial variation in AMR and related traits in the mPact strain panel, representative of the genomic diversity encountered in the opportunistic, high priority human pathogen *P. aeruginosa*, which is a member of the so-called ESKAPE pathogens and which the WHO considers to be a high priority candidate for the development of new treatment options (5). In detail, we identified consistent variation in resistance against a range of clinically relevant antibiotics. Importantly, we also found variation in traits indicating additional responses to antibiotics, including the presence of microcolonies and time-kill dynamics under high drug concentrations. The variation in resistance could be linked to the presence of known AMR genes in the included strains, which are usually found on mobile genetic elements, especially ICE/IMEs or plasmids. Surprisingly, we could not infer strong correlations between the distinct resistance phenotypes and also not between these and the described general growth characteristics. The only exception is the observed relationship between ceftazidime resistance and ceftazidime time-kill dynamics. Moreover, we developed a new approach that integrates data on resistance, growth, virulence, and life-history characteristics and that demonstrates the presence of distinct adaptive strategies among the mPact strains. Overall, we here provide a reference data set for a strain panel that is representative of the genomic diversity of the species *P. aeruginosa*, covering different environmental as well as clinical origins. The mPact strain panel is of a manageable size and clearly smaller than alternative *P. aeruginosa* strain panels (25–27), thus facilitating in-depth molecular as well as evolutionary analyses in the future that go beyond the two main laboratory strains PAO1 and PA14.

The results of our different antimicrobial resistance measurements correlated strongly, indicating that phenotypic resistance is robustly expressed in liquid and agar culture as well as minimal medium. Minor limitations to this agreement were observed for some beta-lactam antibiotics and colistin. Automated susceptibility testing evaluated four strains to be resistant to piperacillin or ceftazidime, which was not confirmed by the other measurements. However, reduced susceptibility was observed in all cases, which suggests that the issue lies with the resistance breakpoints rather than an inconsistency in resistance expression. The lack of strongly resistant strains against the drugs in the panel exacerbates this problem. Reliable colistin resistance measurements, on the other hand, are notoriously difficult (28). We identified genetic resistance determinants for the observed fluoroquinolone and aminoglycoside resistance. Specifically, point mutations in DNA gyrase and topoisomerase are frequently encountered in quinolone-resistant strains of many species, including *P. aeruginosa* (29). Different aminoglycoside-modifying enzymes (AMEs) were found in the two highly aminoglycoside-resistant strains H03 and H06. In contrast, no established beta-lactam resistance genes were found, although our manual analysis found a frameshift mutation in dacB (penicillin-binding protein 4), which is involved in cell wall recycling and strongly induces expression of the chromosomal beta-lactamase AmpC (blaPDC) when inactivated by mutation (30). In general, beta-lactam resistance in *P. aeruginosa* can result from a multitude of mechanisms including porin loss, efflux pumps, beta-lactamases, and target modification, making reliable prediction difficult (31).

The resistance phenotypes were found to be stably expressed in our drug-free evolution experiments, with one notable exception. Strain H01 exhibited strong loss of beta-lactam and especially aminoglycoside resistance. While we found no distinct beta-lactam resistance genes, a unique AME was identified in ant(2’’)-Ia (synonym *aadB*). This aminoglycoside adenylyltransferase was previously reported on mobile genetic elements in *P. aeruginosa* and other Gram-negative pathogens (32). This loss of resistance suggests high costs to the cell, resulting in rapid loss of the phenotype once the selective pressure is lifted. In many cases, this is associated with location of the resistance genes on a costly plasmid, which may get lost after removal of the selective constraint. This scenario cannot explain our finding, since H01 does not seem to bear a plasmid and, thus, the exact underlying reasons still remain to be determined in future work. Irrespective of the exact cause, such rapid resistance losses may have implications for clinical diagnostics and experimentation where the studied strains are repeatedly precultured without drugs, favoring lineages with low copy number or even loss of the entire mechanism (33–35). Overall, the observed resistances did not coincide with phylogenetic relatedness. Instead, the most resistant strains had a clinical background, suggesting that individual strain histories influence their resistances (although a few environmental strains also produced moderate levels of resistance). This general pattern is supported by our reconstruction of genetic determinants for surface disinfectant resistance in the resistant clinical strains.

We additionally examined the strains for the presence of microcolonies in the zones of inhibition and their effect on MICs. This phenomenon indicates that a bacterial population, despite being monogenically founded, has members with increased resistance. When resulting in extensive increases in the overall resistance of the population, this is called heteroresistance, and can have strong implications for diagnostics. For one, microcolonies may not always be visible for inspection after the regular 18-24 hour incubation, leading to underestimation of MICs and potentially misclassification as susceptible. Further, heteroresistance is a possible stepping stone towards full resistance, meaning the population may be in the process of losing its susceptibility. The exact clinical implications of this phenomenon are not fully understood. Unexpectedly, we found comprehensive variation among strains and drugs, suggesting a variety of underlying mechanisms rather than a drug-specific effect or a strain-specific mechanism, such as increased mutation rates (36). Further investigation of these phenotypes and their consequences for adaptation to antimicrobials is warranted.

Variation in strain responses to antibiotics was also found in time-kill kinetics. After exposing the panel strains to above-MIC drug concentrations, the decrease in cell density occurred at different speeds. Generally, slow decrease in cell number could be an indication that strains handle antimicrobial stress through tolerance rather than phenotypic resistance (37). We observed a pattern of drug-specific differences in time-kill kinetics, with the beta-lactams ceftazidime and meropenem generally reducing cell density more slowly than ciprofloxacin or gentamicin. This is likely at least in part the result of their mechanism of action. Beta-lactam antibiotics inhibit peptidoglycan synthesis, affecting both cell wall maintenance and cell division. Therefore, the speed of bactericidal action may depend on the growth speed and conditions. Pharmacodynamic parameters may also play a role in these kinetics, because beta-lactams primarily bind their targets covalently, meaning that their effective concentration decreases over time. This effect is also the basis for the so-called inoculum effect (38). We further observed a correlation of ceftazidime MIC and area under the time-kill curve, suggesting that more resistant strains are killed more slowly, despite the fact that all strains were treated with at least 32x their MIC. In contrast, no correlation was found between growth characteristics and time-kill dynamics, indicating that tolerance by generally slow growth does not play a role in these observations.

We further explored the genomic potential of the mPact panel by employing metabolic and ecological modeling. We used the available whole genome sequences to infer life history characteristics and predict phenotypes of the strains. On the one hand, we found stress-tolerance traits consistently associated with clinical isolation source, higher virulence, and faster growth rate. Interestingly, effector delivery systems such as secretion systems, potentially involved in antibiotic resistance, were found to be predictive for virulent strains. On the other hand, carbohydrate degradation was prominent for environmentally isolated strains and lower virulence. In recent years, life history traits analysis has been applied to analyze environmental and antibiotic-resistant microorganisms (39, 40). In particular, stress-response such as resistance to antibiotics are known to impose a burden on microbial physiology with consequences for growth and resource acquisition (41, 42). Although we did not see an impact of stress-response on growth rate, we found less capacity for carbohydrate acquisition for virulent and clinical strains. Higher growth rates were associated with the enrichment of genes involved in heat response and host interactions, which might link to host adaptation and strategy specific for pathogens (43, 44). In addition, investment in versatile carbohydrate degradation pathways can negatively impact growth (45) and shapes the life history strategy for many environmental microbes (46). Therefore, we hypothesize that mPact strains follow distinct adaptive strategies characterized by either increased degradation or stress and host response. Moreover, we found an opposing pattern in polyamine metabolism. Environmental strains showed an increased capacity to degrade polyamine, and clinical strains showed a higher capacity towards biosynthesis. Polyamines, such as putrescine and spermidine, are ubiquitously present in all organisms, play a prominent role in many processes like gene regulation, cell proliferation, stress response, and can affect host lifespan (47–49). Several host-associated microbes can produce polyamines (50) and especially pathogens are known to interfere with host polyamine metabolism (51, 52). Therefore, the opposing pattern of polyamine degradation and biosynthesis further supports our proposed classification into degradation and host-related strategies for the mPact strains. To our knowledge, this is the first application of life history theory to microorganisms on the strain level.

Altogether, we here present a panel of 22 strains of *P. aeruginosa*, representing the species’ genomic diversity and a variety of resistance phenotypes and growth characteristics. The mPact panel is of suitable size for the in-depth analysis of any phenotypes of interest, without constraining the results to the usual lab strains and therefore greatly expanding the predictability of general application. Most strains are highly susceptible to a range of clinically relevant antibiotics, making them suitable for the examination of the trajectories of resistance evolution of variable genetic backgrounds using experimental evolution. All but one genome have been fully sequenced and are publicly available. Life histories analysis suggests differences in strain adaptation dependent on isolation source or virulence. The mPact panel is therefore a useful resource for fundamental and translational research of this important human pathogen.

## Materials and Methods

Experiments were conducted with a strain panel of *Pseudomonas aeruginosa* which represents genomic diversity observed in the species. Details on strain origin, microassay classification and whole genome sequencing has previously been published (8, 20, 22). The reference strains PA14 (18, 19) and PAO1 (17) were also included. Bacteria were grown in M9 minimal medium supplemented with glucose (2 g/L), citrate (0.58 g/L), and casamino acids (1 g/L) or on M9 minimal agar (1.5%), unless otherwise indicated. Antibiotics were added as specified in the respective sections. Cultures and plates were incubated at 37°C. Statistical analysis was performed in R, Version 4.3.1. All R scripts are included in the supplement.

### Phylogenomics and identification of mobile genetic elements

Phylogenetic trees were constructed as described in Botelho et al (8). In brief, a total of 5468 *P. aeruginosa* genomes were downloaded from RefSeq’s NCBI database using PanACoTA v1.2.0 (53). After removal of low-quality assemblies, duplicates and misclassified assemblies, 1991 genomes were retained. The genomes of 19 mPact strains were added to this, resulting in a total of 2010 genomes. A maximum likelihood tree was inferred with the General Time Reversible model of nucleotide substitution in IQ-TREE v2.1.2. (54) and visualized with iTOL v6 (55). To search for ICEs/IMEs in the mPact strains genomes, we used the annotated files generated by prokka as input in the standalone-version of ICEfinder (56).

### Antimicrobial susceptibility

The minimum inhibitory concentration of antibiotics was measured using (i) a gradient test strip (57) and (ii) the disc diffusion test (58). Bacterial cultures were incubated at 37°C for 18h, with subsequent inoculation of M9 plates with a cotton swab of overnight culture (OD_600_ at 0.08). (i) MIC test strips (Liofilchem®) were placed onto the plate and the plate was incubated for 24h, after which the MIC was read at the intersection of the zone of inhibition and the test strip. (ii) 3 antimicrobial discs (Mast Diagnostica) per plate were applied with a disc dispenser (MDD65, Mast Diagnostica) and incubated for 24h, after which the inhibition zones were analyzed by standardized image capture using a flatbed scanner (Epson Perfection V600 pro). Image analysis was performed with the program Antibiogram J (59) (always in triplicate). We also performed automated susceptibility testing using the VITEK2 system (bioMérieux) and analyzed using the EUCAST clinical breakpoint table Version 13.1 (60). Breakpoints for gentamicin were taken from Version 9, the last version to contain values for this drug in *P. aeruginosa* (61).

### Presence of microcolonies

MIC test strip plates were visually inspected for the presence of microcolonies inside the zones of inhibition. If present, MICs with and without these colonies were recorded. Plates were incubated for another 24 hours to improve colony detection. No differences were observed between the two time points (data not shown).

### Assessment of resistance stability

Resistance stability was assessed after evolution in the absence of drug. Populations of 3 independent parallel cultures per strain were seeded in antibiotic free medium and grown for 18h. They were then diluted to 0.08 and 100 uL was plated onto plates without antibiotics. Every 24h, 20% of the bacteria were transferred to a fresh plate without antibiotic. The experiment lasted a total of 15 days. Evolved populations were frozen in dimethyl sulfoxide on the last day. Due to logistic reasons, only 18 out of 22 strains of the panel were used and had to be split in two separate evolution experiments. As controls, two strains were included in both. Results for them were found to be consistent across the experiments (Kolmogorov-Smirnov test). Resistance stability was calculated as log2-transformed means of replicates minus log2-transformed ancestral MICs.

### Time-kill curves

Measures for antibiotic tolerance were based on time-kill curves (37). In brief, each strain was cultured in M9 medium overnight from single colonies and the resulting culture was adjusted to OD_600_ 0.08 - 0.12. After that, antibiotics (at least 10 times higher than MIC, Meropenem: 15 μg/ml, Gentamycin: 30 μg/ml, Ciprofloxacin: 15 μg/ml, Ceftazidime: 200 μg/ml) were added to the diluted bacterial culture and the tubes incubated at 37℃ with shaking. At different time points, the number of cells was monitored. Serial dilution and plating were used to determine the number of colony forming units. The survival frequency was calculated by the number of colonies in different time points divided by that of starting point. To exclude phenotypic resistance effects on the time-kill curve analysis, strains where the effective concentrations were less than 32x the MIC were excluded from analysis. For all other combinations, the area under the curve was calculated using the AUC function of the DescTools package in R with the trapezoid approach.

### Growth curves

Growth curves were assessed in 96-well plates. In short, we inoculated one colony overnight for 20h in LB-Medium at 37 °C with shaking. 0.5% of the overnight liquid culture were subsequently inoculated in the 96-well plates in a fully randomized way containing M9 medium. The growth kinetics were assessed with continuous shaking (Shake mode: double orbital; Orbital frequency: 807 cpm (1mm); Orbital speed: Fast), and optical density measurements at 600nm in a plate reader (Epoch 2, Agilent) for 20h, with 15 min time points. For each strain three biological replicates were tested. Growth parameters were modeled using the method by Hall et al. (24), implemented in the growthrates package in R (easylinear method), yielding maximum growth rates and length of lag phase. Replicates with model fit scores <0.9 were excluded. Carrying capacity was extracted from the OD kinetic over time. AUC was calculated as mentioned above, using a spline approach.

### Antimicrobial resistance genes

Genomic resistance genes were retrieved by parsing the assembled and closed genomes into AMRFinderPlus(62). Hits encountered in all strains were filtered manually. For additional resistance genes, the amino acid sequences of the genes in question were extracted from the genomes’ RefSeq annotations by downloading the gbff files from Genbank, splitting the contigs into individual files, and loading them into a common dataset using the genbankr package in R with bioconductor 3.17 (63). Coding sequences were extracted with the GenomicFeatures package and aligned using the msa command with default settings. Neighbor-joining trees were generated with the nj command of the ‘ape’ package and plotted using tidytree.

### Metabolic models and life history

Metabolic models were reconstructed from mPact genomes by employing Bactabolize v1.02 (64), and the enrichment analysis was performed using clusterProfiler v4.10 (65). Life history traits were inferred by eggNOG-Mapper v2.1.12 (DB v5.0.2) (66), bakta v1.9 (DB 5.0) (67), dbCAN v4.0 (68), abricate v.1.0.1 (DB vfdb 2023-Aug-18) (69, 70), antiSMASH 6.1.1, gRodon v.2.3.0 (71), and MetaCyc metabolic pathways (72) were predicted by gapseq v1.2 (73). R v4.3.1 was used for the statistical analysis, and associated life history traits were selected by Boruta v8.0 (74) and glmnet v4.1.7. The predictive quality was accessed using caret v6.0-94 (leave-one-out cross-validation, tuneLength=1000) (75).

## Acknowledgements

We thank Tabea Loeblein, Fernanda Fontes Trancoso, Konstantin Kempe, Laura Kirchhoff, and Kira Haas for supporting the experimental work and the Rupp and Schulenburg groups for general feedback. We are grateful for financial support from the German Science Foundation within the Research and Training Group 2501 (RTG 2501) on Translational Evolutionary Research (project 4.2 to HS), within the Excellence cluster Precision Medicine in chronic Inflammation (PMI; funding under Germany’s Excellence Strategy EXC 2167-390884018, to JR, HS), and as part of the individual grant SCHU 1415/12-2 (to HS). We are also grateful for financial support from the Max-Planck Society (Fellowship to HS), the Leibniz Association within the Leibniz Science-Campus Evolutionary Medicine of the Lung (EvoLUNG, to HS), and the Chinese Scholarship Council (to JL). This research was supported in part through high-performance computing resources available at the Kiel University Computing Centre. The funders had no role in study design, data collection and interpretation, or the decision to submit the work for publication.

## Transparency declarations

Conceptualisation: LT, AB, JZ, JR, FB, HS

Formal analysis: LT, AB, JZ, FB, JL, AM, JK, NMM

Funding acquisition: HS, JR, BT

Investigation and methodology: LT, AB, JZ, FB, JL, JB, NMM

Resources: AM, JK, BT

Supervision: LT, AB, JZ, FB, JR, BT, HS

Writing of original draft: LT, AB; JZ, FB; JL, HS

Writing: review and editing: all authors.

## Supplement

**Figure S1:**
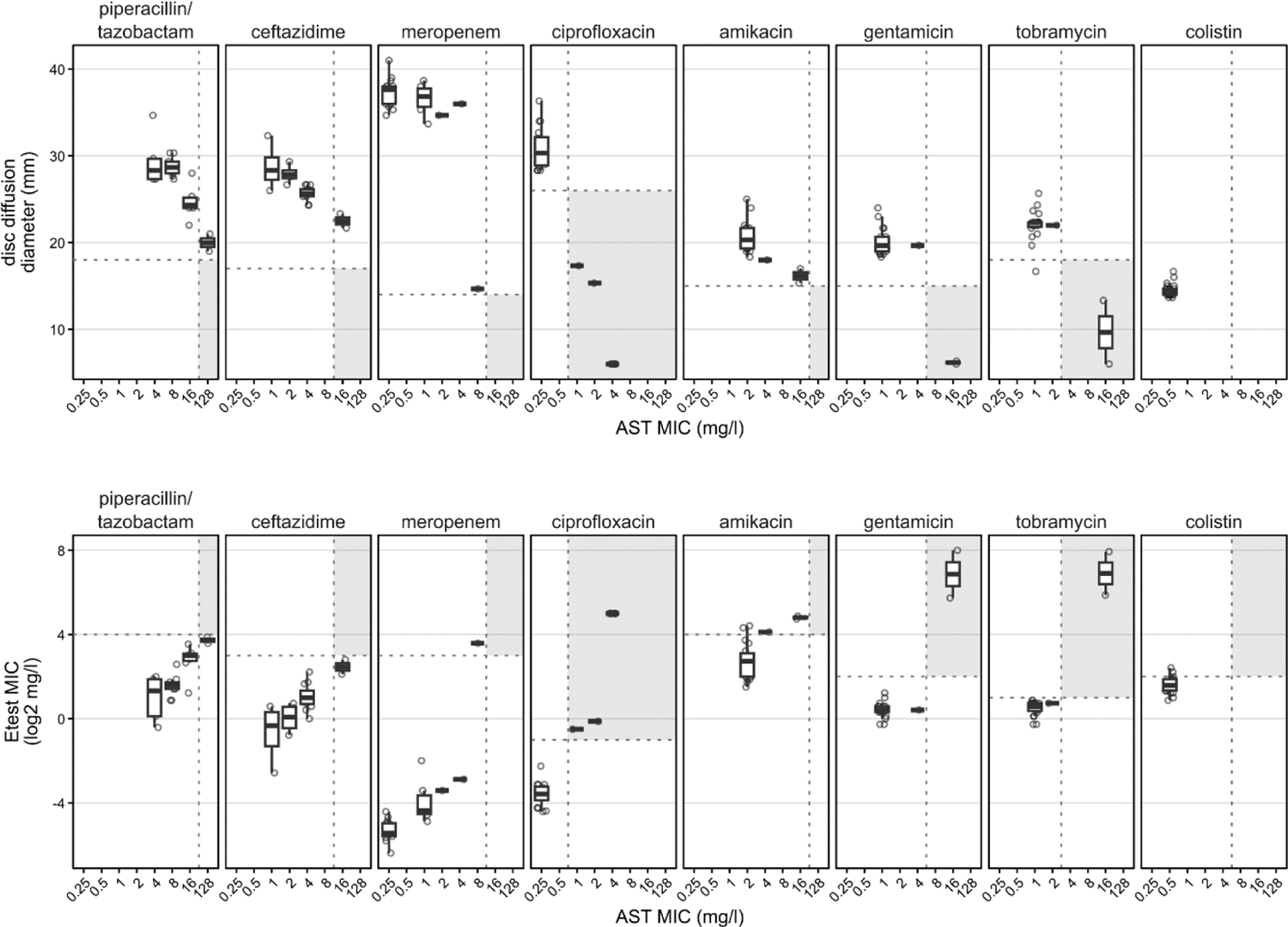
Comparison of automated susceptibility testing and M9 agar methods Resistance breakpoints per method are indicated by dashed lines (AST vertical, M9 methods horizontal. Zones of categorical agreement of classification as ‘resistant’ are marked in gray.

**Figure S2:**
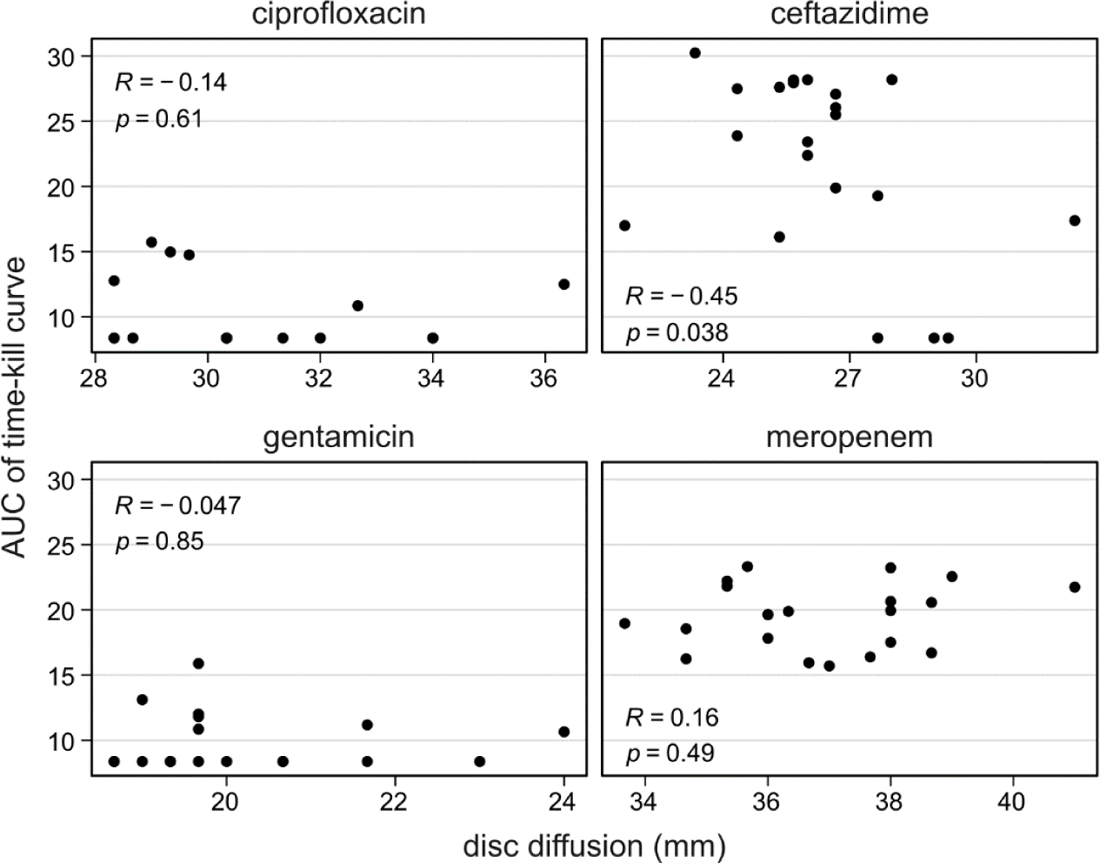
Correlation of strain MIC and area under the time-kill curve Coefficient of correlation (R) and *p*-values of the t-test of *R* = 0.

**Figure S3:**
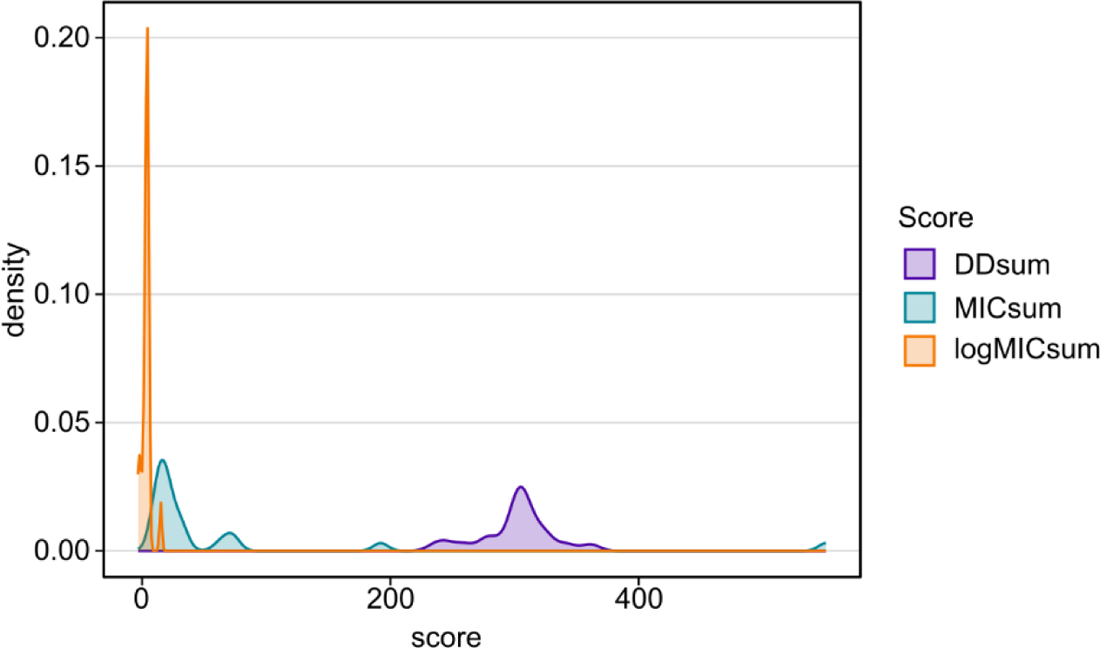
Density of strains’ cumulative resistance scores. Sum of disc diffusion diameters (DDsum), sum of gradient strip MICs (MICsum) and log2 transformed sum of differences between MICs and drug resistance breakpoints (logMICsum). Details see text.

**Figure S4:**
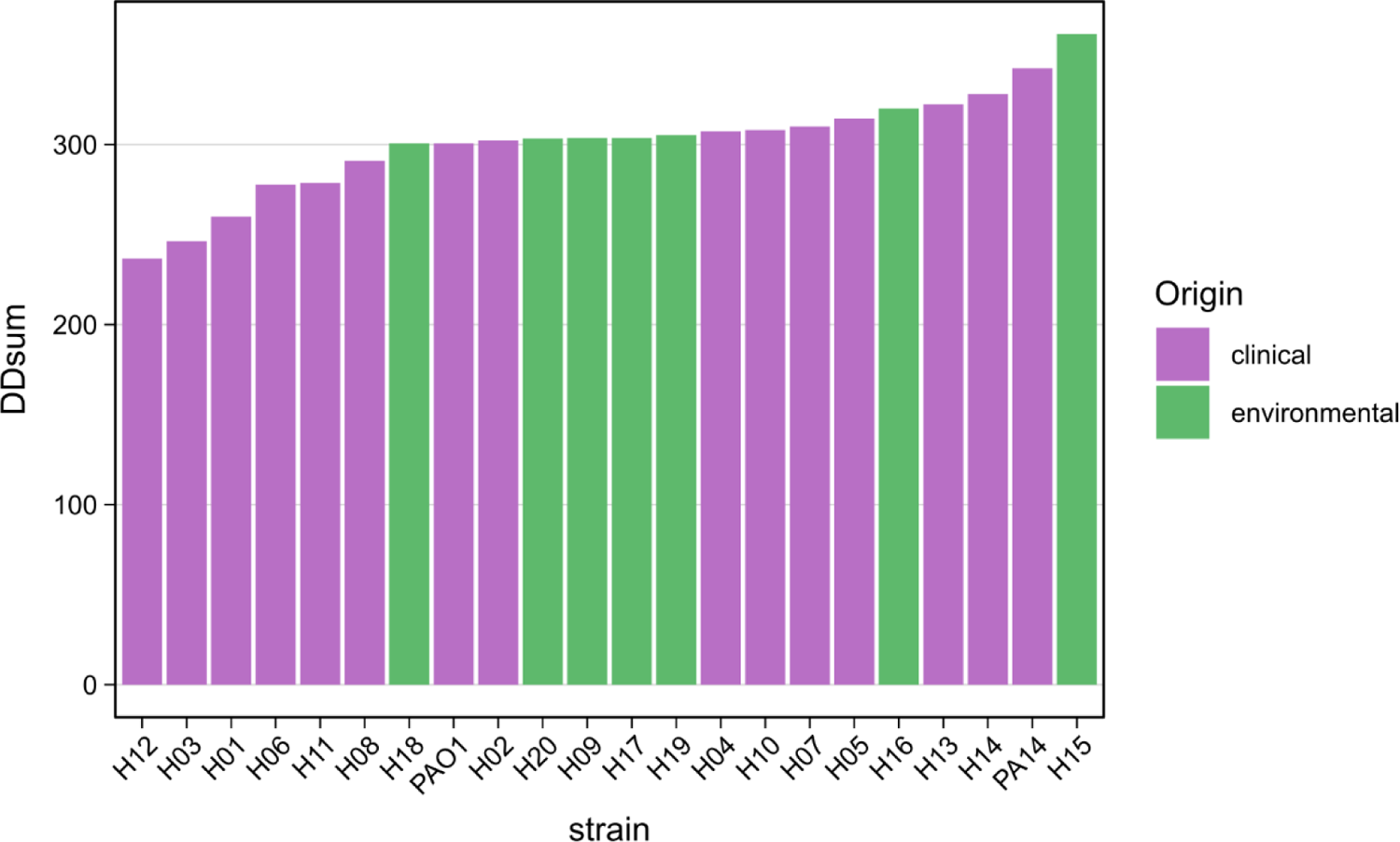
Strains sorted by cumulative resistance. Sum of disc diffusion diameters (DDsum) with strain origin marked as color.

**Figure S5:**
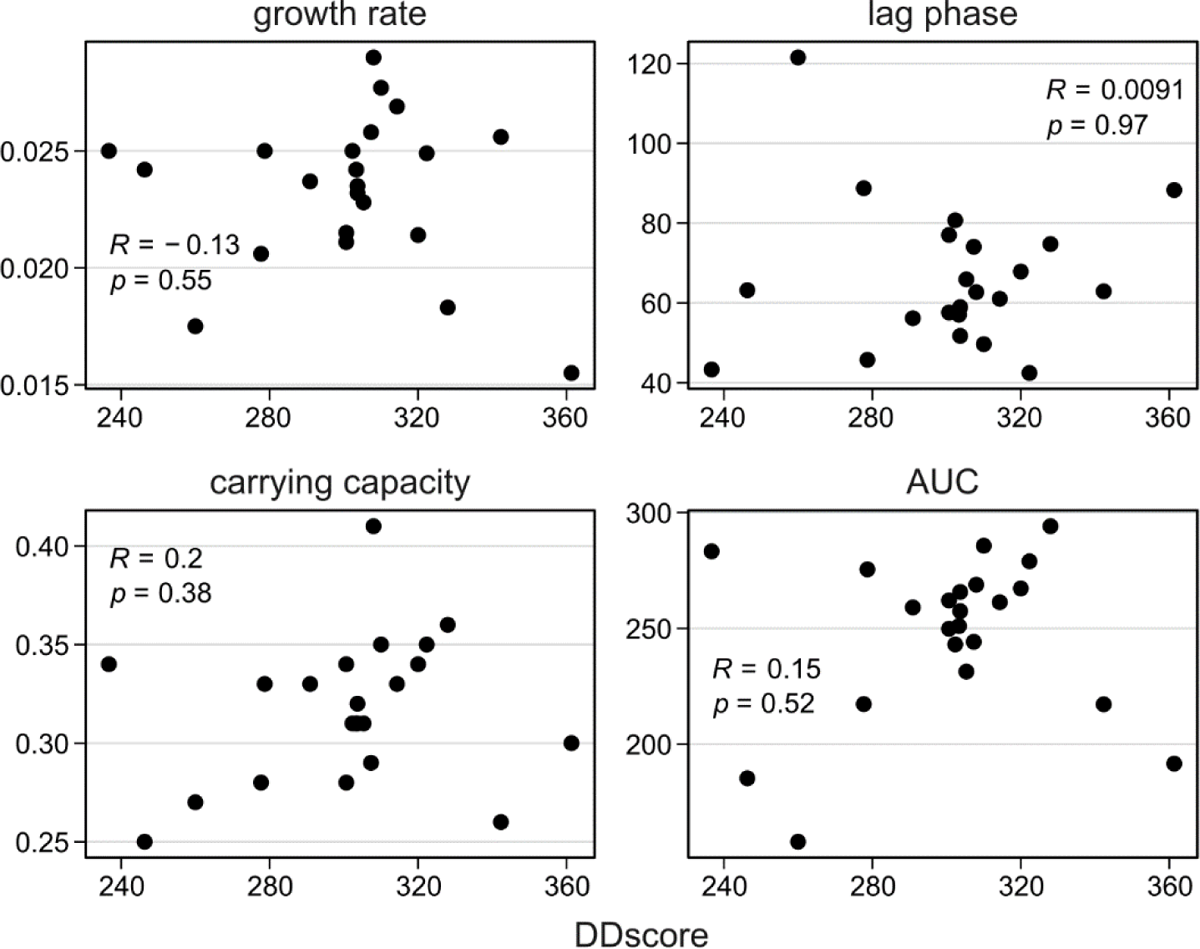
Correlation of growth parameters and resistance. The sum of disc diffusion diameters for x drugs (DDScore) is correlated with the modeled growth parameters of the individual strains (coefficient of correlation R and t-test for R = 0 as p)

**Figure S6:**
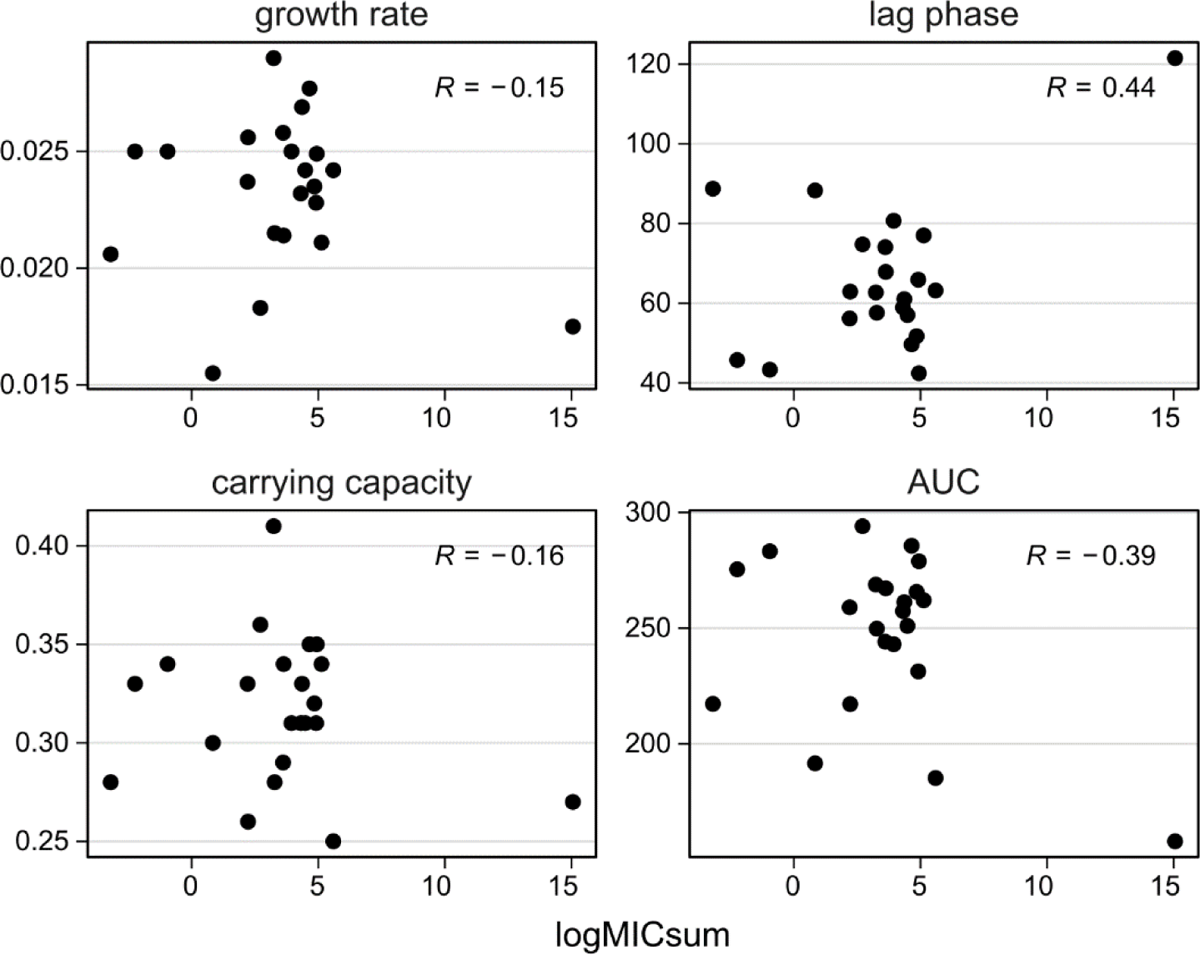
Correlation of growth parameters and gradient strip resistance. The log2 transformed sum of differences between MICs and drug resistance breakpoints (logMICsum) is correlated with the modeled growth parameters of the individual strains.

**Figure S7:**
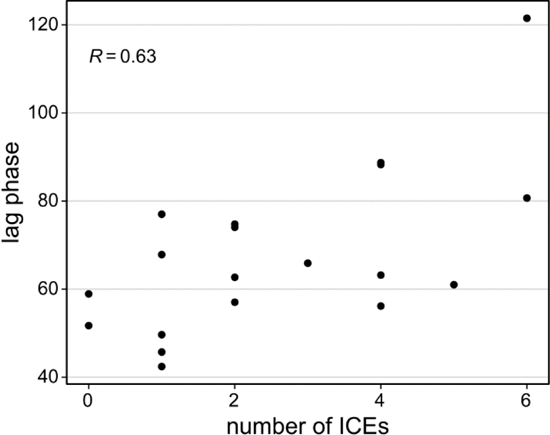
Correlation of lag phase and number of integrative conjugative elements per strain

**Figure S8:**
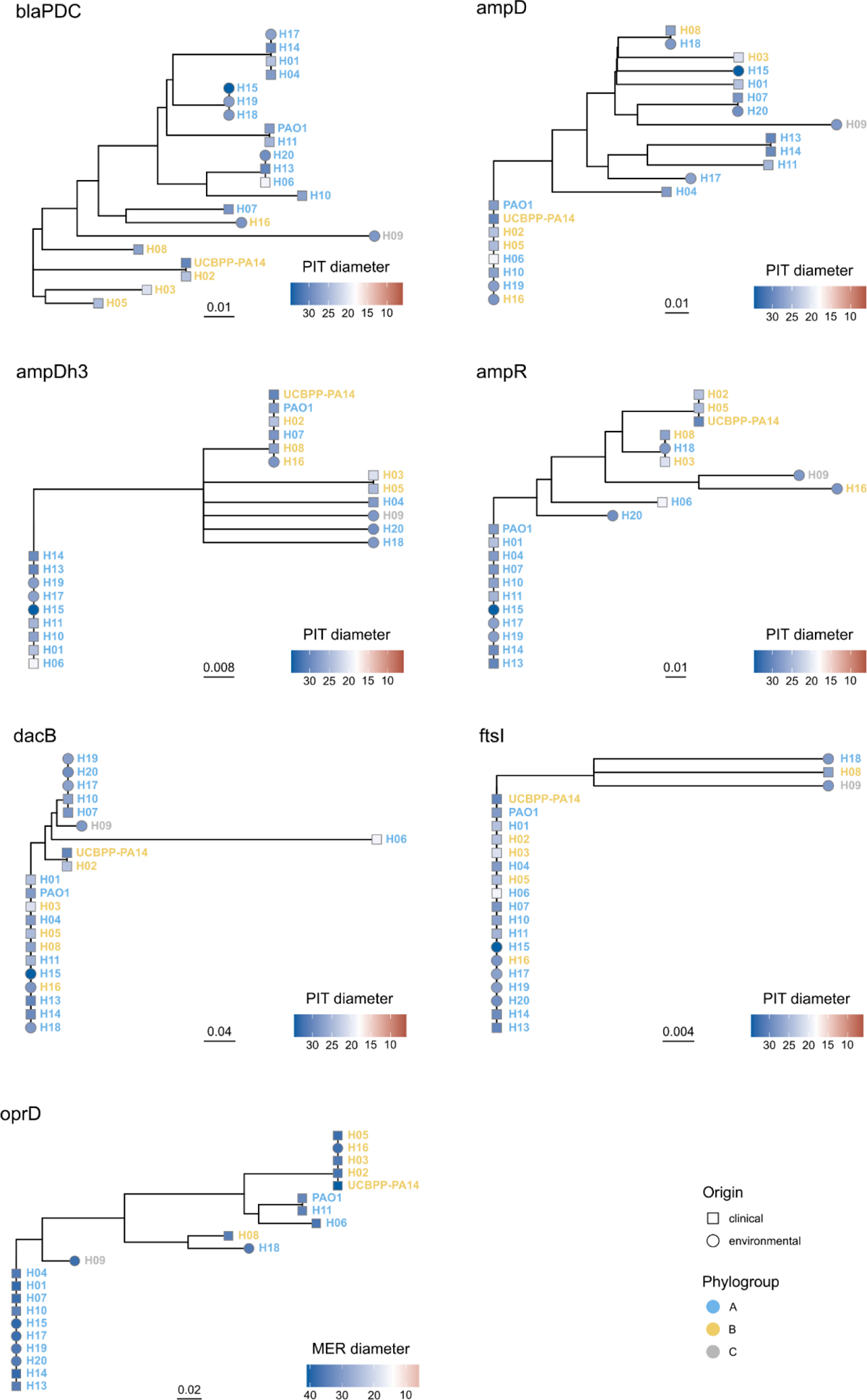
Betalactam resistance gene phylogenies. Neighbor-joining trees of amino acid sequences of the indicated genes. Strain names at the edges are shaded by phylogroup and shaped by strain origin. Resistance to an indicator drug (PIT = piperacillin/tazobactam, MER = meropenem) is given as the color of the edges, with the values centered around the resistance breakpoints in white and values below in blue.

**Table S1:** Modeled growth parameters.

| Strain | Growth rate (mean) | Growth rate (sd) | Lag phase (mean) | Lag phase (sd) | Carrying capacity (mean) | Carrying capacity (sd) | AUC (mean) | AUC (sd) |
| --- | --- | --- | --- | --- | --- | --- | --- | --- |
| H01 | 0.0175 | 0.0053 | 121.51 | 52.31 | 0.27 | 0.03 | 157.84 | 19.55 |
| H02 | 0.025 | 0.007 | 80.69 | 35.32 | 0.31 | 0.04 | 243.05 | 23.54 |
| H03 | 0.0242 | 0.0049 | 63.18 | 27.93 | 0.25 | 0.02 | 185.23 | 22.2 |
| H04 | 0.0258 | 0.0071 | 74.05 | 43.51 | 0.29 | 0.04 | 244.16 | 26.43 |
| H05 | 0.0269 | 0.0058 | 61.01 | 20.47 | 0.33 | 0.02 | 261.26 | 9.38 |
| H06 | 0.0206 | 0.0064 | 88.73 | 33.39 | 0.28 | 0.02 | 217.26 | 19.79 |
| H07 | 0.0277 | 0.0057 | 49.65 | 20.54 | 0.35 | 0.05 | 285.66 | 25.95 |
| H08 | 0.0237 | 0.0058 | 56.16 | 27.71 | 0.33 | 0.08 | 259.02 | 29.88 |
| H09 | 0.0232 | 0.0044 | 58.92 | 16.81 | 0.31 | 0.03 | 257.41 | 33.68 |
| H10 | 0.029 | 0.0036 | 62.69 | 13.55 | 0.41 | 0.06 | 268.85 | 21.24 |
| H11 | 0.025 | 0.0068 | 45.72 | 9.98 | 0.33 | 0.05 | 275.44 | 28.74 |
| H12 | 0.025 | 0.0072 | 43.29 | 18.82 | 0.34 | 0.05 | 283.25 | 26.85 |
| H13 | 0.0249 | 0.0037 | 42.42 | 30.91 | 0.35 | 0.04 | 278.95 | 18.58 |
| H14 | 0.0183 | 0.0056 | 74.76 | 42.92 | 0.36 | 0.04 | 294.11 | 23.02 |
| H15 | 0.0155 | 0.0036 | 88.28 | 20.78 | 0.3 | 0.04 | 191.56 | 21.3 |
| H16 | 0.0214 | 0.0042 | 67.84 | 24.05 | 0.34 | 0.04 | 267.23 | 11.79 |
| H17 | 0.0235 | 0.0059 | 51.72 | 16.97 | 0.32 | 0.04 | 265.71 | 32.13 |
| H18 | 0.0211 | 0.0047 | 77.01 | 30.51 | 0.34 | 0.04 | 262.06 | 19.73 |
| H19 | 0.0228 | 0.0061 | 65.89 | 25.73 | 0.31 | 0.04 | 231.31 | 27.21 |
| H20 | 0.0242 | 0.0057 | 57.03 | 20.78 | 0.31 | 0.02 | 251 | 29.15 |
| PA14 | 0.0256 | 0.006 | 62.9 | 22.06 | 0.26 | 0.02 | 217.18 | 13.47 |
| PAO1 | 0.0215 | 0.0035 | 57.59 | 20.12 | 0.28 | 0.02 | 249.8 | 9.75 |

AUC: area under the growth curve; SD: standard deviation. Lag phase given in minutes. Carrying capacity given as OD.

**Table S2:** Statistical test overview.

| analysis | method | test statistic | fdr-corrected *p* |
| --- | --- | --- | --- |
| phylogroup A/B - logMICsum | Wilcoxon rank sum exact test | W = 44 | 1 |
| phylogroup ABB - DDsum | Wilcoxon rank sum exact test | W = 41 | 1 |
| origin - logMICsum | Wilcoxon rank sum exact test | W = 35 | 0.75 |
| origin - DDsum | Wilcoxon rank sum test with continuity correction | W = 40 | 0.75 |
| plasmids - logMICsum | Wilcoxon rank sum exact test | W = 2 | 0.94 |
| plasmids - DDsum | Wilcoxon rank sum test with continuity correction | W = 5 | 1 |
| logMICsum ~ ICEs | Pearson's product-moment correlation | t = 0.97 | 0.75 |
| DDsum ~ ICEs | Pearson's product-moment correlation | t = -1.105 | 0.75 |
| origin - growthrate | Wilcoxon rank sum test with continuity correction | W = 81.5 | 0.31 |
| origin - lagphase | Wilcoxon rank sum exact test | W = 45 | 0.941 |
| origin - auc | Wilcoxon rank sum exact test | W = 58 | 0.97 |
| origin - carryingcapacity | Wilcoxon rank sum test with continuity correction | W = 51.5 | 1 |
| phylogroup A/B - growthrate | Wilcoxon rank sum test with continuity correction | W = 29 | 0.75 |
| phylogroup A/B - lagphase | Wilcoxon rank sum exact test | W = 41 | 1 |
| phylogroup A/B - auc | Wilcoxon rank sum exact test | W = 54 | 0.75 |
| phylogroup A/B - carryingcapacity | Wilcoxon rank sum test with continuity correction | W = 51.5 | 0.782 |
| plasmids - growthrate | Wilcoxon rank sum test with continuity correction | W = 30 | 0.75 |
| plasmids - lagphase | Wilcoxon rank sum exact test | W = 11 | 0.75 |
| plasmids - auc | Wilcoxon rank sum exact test | W = 27 | 0.94 |
| plasmids - carryingcapacity | Wilcoxon rank sum test with continuity correction | W = 22.5 | 1 |
| ICEs ~ growthrate | Pearson's product-moment correlation | t = -0.96 | 0.75 |
| **ICEs ~ lagphase** | **Pearson's product-moment correlation** | **t = 3.366** | **0.044** |
| **ICEs ~ auc** | **Pearson's product-moment correlation** | **t = -3.6** | **0.04** |
| ICEs ~ carryingcapacity | Pearson's product-moment correlation | t = -2.095 | 0.309 |

